# Towards chemical accuracy for alchemical free energy calculations with hybrid physics-based machine learning / molecular mechanics potentials

**DOI:** 10.1101/2020.07.29.227959

**Authors:** Dominic A. Rufa, Hannah E. Bruce Macdonald, Josh Fass, Marcus Wieder, Patrick B. Grinaway, Adrian E. Roitberg, Olexandr Isayev, John D. Chodera

## Abstract

Alchemical free energy methods with molecular mechanics (MM) force fields are now widely used in the prioritization of small molecules for synthesis in structure-enabled drug discovery projects because of their ability to deliver 1–2 kcal mol^−1^ accuracy in well-behaved protein-ligand systems. Surpassing this accuracy limit would significantly reduce the number of compounds that must be synthesized to achieve desired potencies and selectivities in drug design campaigns. However, MM force fields pose a challenge to achieving higher accuracy due to their inability to capture the intricate atomic interactions of the physical systems they model. A major limitation is the accuracy with which ligand intramolecular energetics—especially torsions—can be modeled, as poor modeling of torsional profiles and coupling with other valence degrees of freedom can have a significant impact on binding free energies. Here, we demonstrate how a new generation of hybrid machine learning / molecular mechanics (ML/MM) potentials can deliver significant accuracy improvements in modeling protein-ligand binding affinities. Using a nonequilibrium perturbation approach, we can correct a standard, GPU-accelerated MM alchemical free energy calculation in a simple post-processing step to efficiently recover ML/MM free energies and deliver a significant accuracy improvement with small additional computational effort. To demonstrate the utility of ML/MM free energy calculations, we apply this approach to a benchmark system for predicting kinase:inhibitor binding affinities—a congeneric ligand series for non-receptor tyrosine kinase TYK2 (Tyk2)—wherein state-of-the-art MM free energy calculations (with OPLS2.1) achieve inaccuracies of 0.93±0.12 kcal mol^−1^ in predicting absolute binding free energies. Applying an ML/MM hybrid potential based on the ANI2x ML model and AMBER14SB/TIP3P with the OpenFF 1.0.0 (“Parsley”) small molecule force field as an MM model, we show that it is possible to significantly reduce the error in absolute binding free energies from 0.97 [95% CI: 0.68, 1.21] kcal mol^−1^ (MM) to 0.47 [95% CI: 0.31, 0.63] kcal mol^−1^ (ML/MM).

## Introduction

### MM force fields are widely used in structure-enabled drug discovery

Alchemical free energy calculations are now widely used in structure-enabled drug discovery programs to optimize or maintain potency [1–5]. Typically, relative alchemical free energy methods can predict affinities with accuracies of 1–2 kcal mol^−1^ in prospective use in well-behaved, structure-enabled programs [3, 6]. While an accuracy of 1 kcal mol^−1^ is already sufficient to greatly reduce the number of compounds that must be synthesized to achieve desired potency gains, the ability to further improve this accuracy to 0.5 kcal mol^−1^ (“chemical accuracy” [7]) would deliver significant benefits at least as large as the improvement achieved by accuracy improvements from 2 kcal mol^−1^ to 1 kcal mol^−1^ for optimization of potency [8, 9] and selectivity [10, 11]. To achieve “chemical accuracy”, improvements are required to the computational model of the protein-ligand system (the force field) while constraining any increase in computational cost to ensure results can be produced on a timescale viable for active drug discovery projects, as alchemical free energy calculations typically require generating tens to hundreds of nanoseconds of simulation data within a few hours [5, 12].

### Surpassing 1 kcal mol^−1^ accuracy requires model improvements that are difficult to generalize

Relative free energy methods almost universally utilize fixed-charge MM force fields to model small, organic, drug-like molecules and interactions with their respective receptors and aqueous environments, such as GAFF [13, 14], CGenFF [15, 16, 16], or OPLS [17]. Importantly, these popular class I [18, 19] MM force fields have well-characterized drawbacks, in part, because they omit a number of important energetic contributions known to limit their ability to achieve chemical accuracy [7, 20, 21]. For example, while moving to more complex electrostatics models which include fixed multipoles and polarizable dipoles [22] are promising, the development of polarizable force fields that broadly deliver accuracy gains has proven challenging [23–25].

Deficiencies in the modeling of torsions that accurately account for local chemical environment is also a difficult challenge for MM force fields [26]. Indeed, many MM force fields recommend refitting torsion potentials directly to quantum chemical calculations for individual molecules in a bespoke manner, a process considered essential to achieving a 1–2 kcal mol^−1^ level of accuracy in binding free energy calculations [1, 27]. Even so, the environment-dependent coupling between torsions and other valence degrees of freedom [7, 20, 21, 28] (including adjacent torsions [26, 29–32]) makes it difficult for this simple refitting approach to accurately capture often significant ligand conformational reorganization effects [33].

### QM/MM offers a parameterization-free alternative, but at significantly increased cost

Another approach attempts to avoid the parameterization issue altogether by modeling the ligand using QM levels of theory, while treating the remaining atomic environment with an MM force field in a hybrid QM/MM potential [34–38]. QM/MM calculations are orders of magnitude more expensive than the equivalent calculation at the MM level, which has led to attempts to speed up these calculations by either using a low level of QM theory, or reducing the number of QM-level evaluations that are performed. QM/MM simulations for ligand binding are yet limited in accuracy due to the low level of QM theory (often semi-empirical or DFT with limited basis sets) that are computationally practical. This has driven the development of methods that attempt to minimize the amount of QM/MM simulation data that must be generated by computing perturbative corrections to MM alchemical free energy calculations [39–41].

### Machine learning (ML) potentials can reproduce QM energies at greatly reduced cost

Recently, quantum machine learning potentials (ML or QML) [42]—such as those based on neural networks like ANI [43]—have seen success in reproducing QM energetics with orders of magnitude less computational cost than the QM methods they aim to reproduce. The ANI-1x neural network potential [43], for example, is able to reproduce DFT-level energies (*ω*B97X functional with 6-31G* basis set) with a 10^6^ speed up. Indeed, the ANI models are so fast and reproduce quantum chemical data so well that recent approaches have integrated them into bespoke torsion refitting schemes as an alternative to costly QM torsion scans [44]. Notably, the recently-developed ANI-2x supports molecular systems including element types C, H, N, O, as well as F, Cl, and S—ideal for applications to receptor-ligand systems as they cover 90% of drug-like molecules. In particular, for the purpose of this study, ANI-2x covers 100% of the ligands included in the Schrödinger benchmark set for alchemical free energy calculations [45].

## Hybrid ML/MM potentials provide improved ligand energetics

While the notion of simulating an entire solvated protein-ligand system with quantum chemical accuracy—in a manner that avoids molecular mechanics parameterization altogether—with orders of magnitude less effort is immediately appealing, several limitations stand in the way to this: First, the number of elements covered by potentials like ANI [49] are so far rather limited, precluding their application to ions and cofactors; Second, while ML models are orders of magnitude faster than any reasonable QM levels of theory, they are currently still orders of magnitude slower than GPU-accelerated MM simulations, though this gap is expected to close rapidly with both software and hardware improvements. Third, ML models have not yet been parameterized on systems that would ensure well-behaved condensed-phase properties, so that MM may yet provide superior results for treating intermolecular interactions in large, extended systems.

However, hybrid ML/MM models—wherein ligand interactions are treated with ML and the environment and ligand-environment interactions with MM (in analogy to QM/MM [34–38])—could provide a convenient and efficient path to improving accuracy by capturing complex molecular interactions that classical force fields fail to do. Recently, Lahey et. al. [52] demonstrated that by using the ANI-1ccx [53] ML potential (a variant of ANI-1x refit to coupled-cluster calculations) to represent intramolecular interactions of small molecule ligands, accurate binding poses and conformational energies could be afforded to the EGFR inhibitor, erlotinib. Notably, they reported significant discrepancies among torsional energy profiles between the MM and ANI-1ccx potentials [52].

Here, we wondered whether incorporating this higher-accuracy ML treatment of small molecule ligand intramolecular energetics within an MM scheme would lead to quantitative improvements in absolute and relative binding free energies. A particularly convenient hybrid ML/MM formulation corresponds to the functional form given in Equation 1 (and visualized in Figure 1) wherein the potential energy function for a environment (receptor and/or solvent)/ligand system takes the form

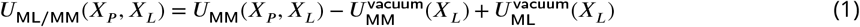

with 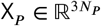 as all non-ligand coordinates (receptor and/or solvent), 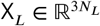 as the ligand coordinates, 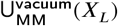 and 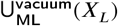 indicating the MM/ML potential energy function for the ligand and U_MM_(X_*P*_, X_*L*_) the MM potential energy function for the environment/ligand system. The formulation given in Equation 1 treats intramolecular ligand interactions with an ML potential while intermolecular and environmental (receptor and/or solvent) atomic interactions are treated with MM force fields [52]. Although other formulations are possible—such as including short-range ligand-environment interactions within the ML region—we will demonstrate that the simple formulation of Equation 1 is sufficient to realize significant improvements in the accuracy of computed binding free energies for a challenging kinase:inhibitor benchmark system (Figure 2).

**Figure 1.**
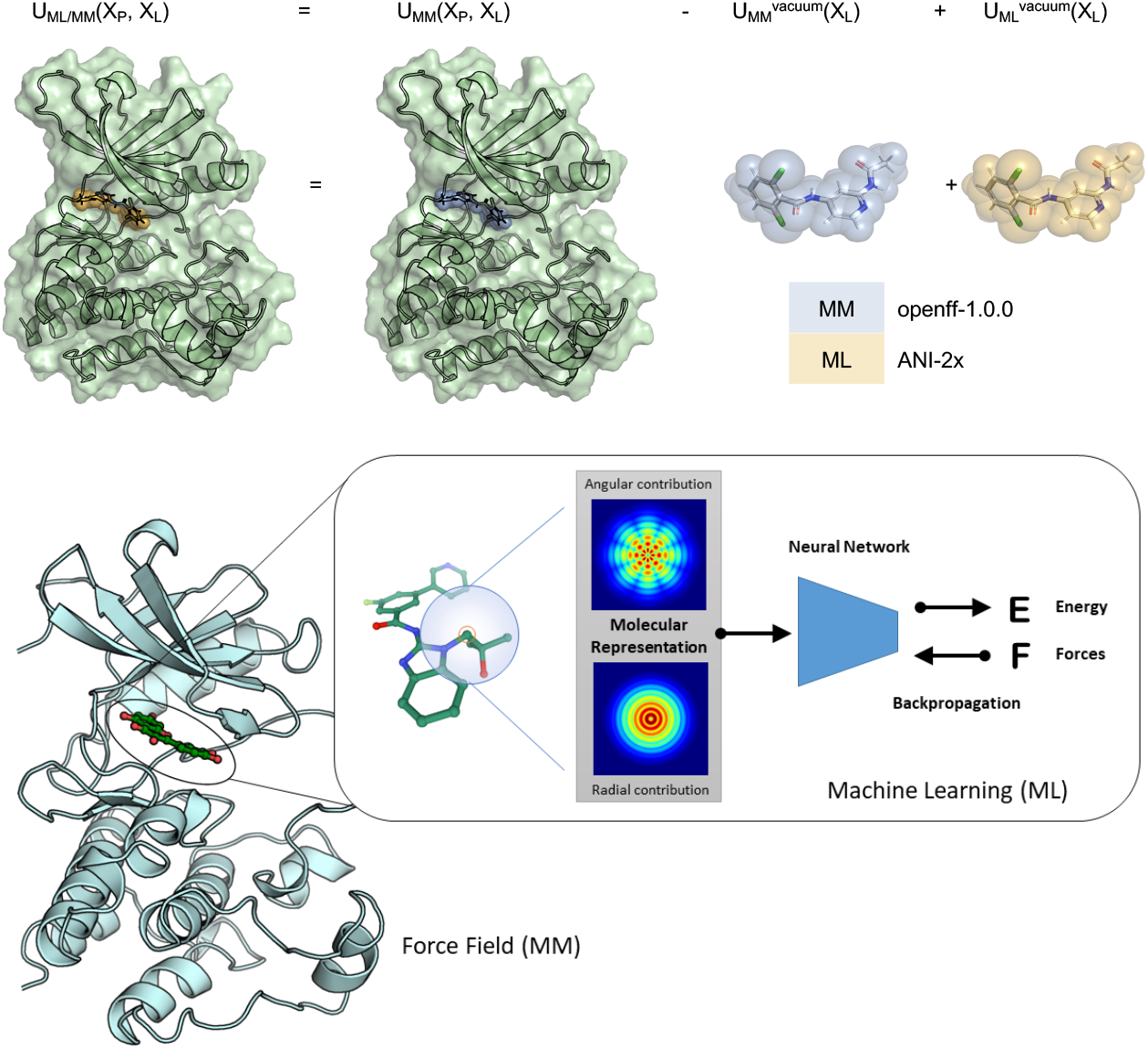
A hybrid ML/MM potential can treat intramolecular ligand forces with high accuracy. ML potentials can treat intramolecular ligand forces with high accuracy, as part of a hybrid ML/MM scheme. *Top:* We can construct a hybrid machine learning / molecular mechanics (ML/MM) potential that treats ligand intramolecular interactions with higher accuracy than achievable by MM potentials by subtracting the MM energy of the ligand in vacuum and adding the more accurate ML energy of the ligand in vacuum. Here, the MM model uses the Open Force Field Initiative [http://openforcefield.org] OpenFF 1.0.0 (“Parsley”) small molecule force field [46], AMBER14SB [47], and TIP3P [48] while the ML model uses the ANI-2x [49] neural network potential parameterized using DFT *ω*B97X/6-31G* QM calculations. *Bottom:* The ANI-2x [49] ML potential first computes radial and angular features for each atom and then sums energetic contributions by atom using deep learning models specific to each element-element pair.

**Figure 2.**
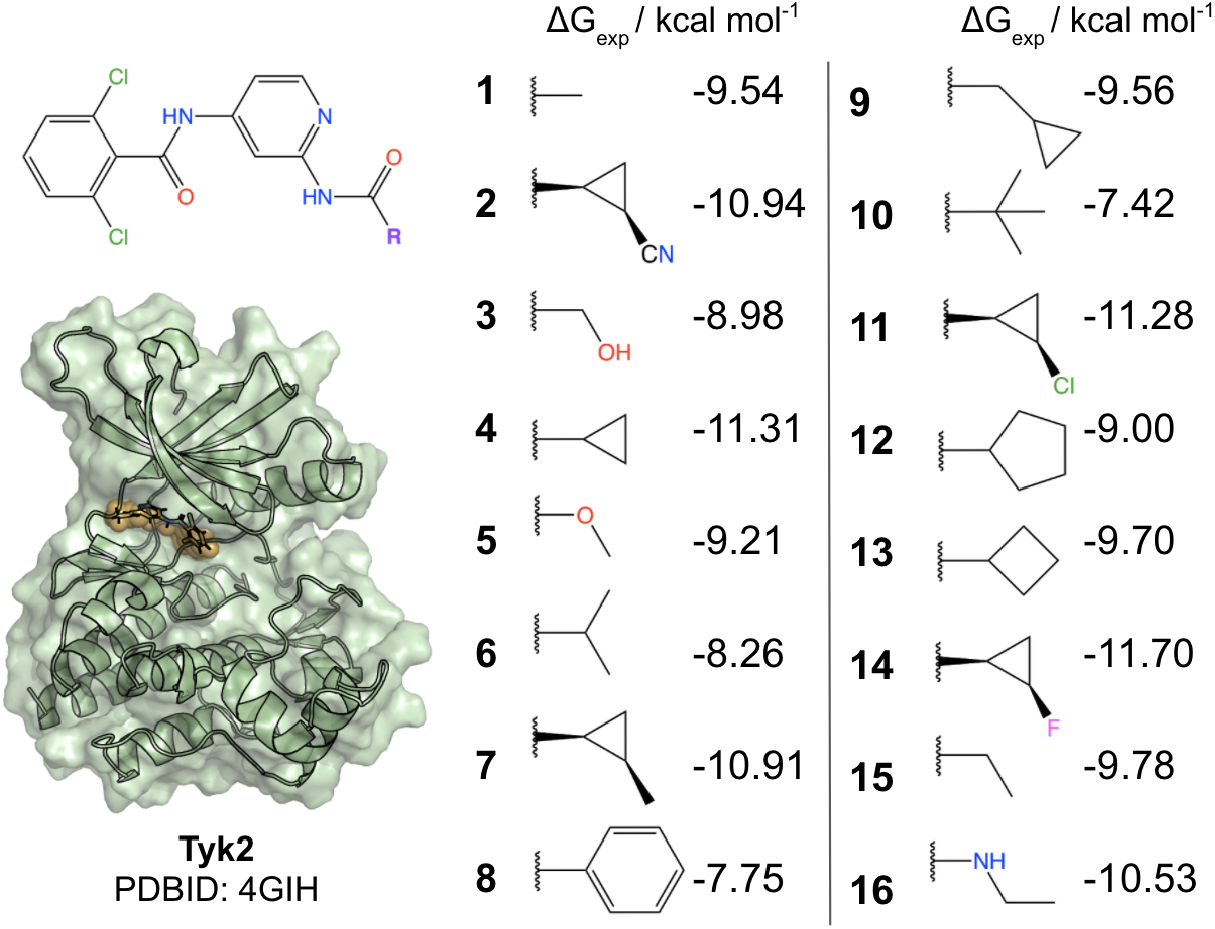
Tyk2 is a challenging test set for predicting kinase:inhibitor binding free energies. The Tyk2 congeneric ligand benchmark series was taken from the Schrödinger JACS benchmark set [45], which is challenging for both commercial force fields (OPLS2.1 achieves a ΔG RMSE of 0.93±0.12 kcal mol^−1^ [45]) and public force fields (GAFF 1.8 achieves a ΔG RMSE of 1.13 kcal mol^−1^, and ΔΔG RMSE of 1.27 kcal mol^−1^ [50]). *Left:* Illustration of the X-ray structure used for all calculations. *Right:* 2D structures of all ligands in the benchmark set, showing common scaffold and substituents. The Schrodinger Tyk2 benchmark set contains a congeneric series selected from [51, 51] where experimental errors in *K_i_* are reported to have *δK_i_*/*K_i_* < 0.3, yielding *δ*Δ*G* ≈ 0.18 kcal mol^−1^ and *δ*ΔΔ*G* ≈ 0.25 kcal mol^−1^.

**Figure 3.**
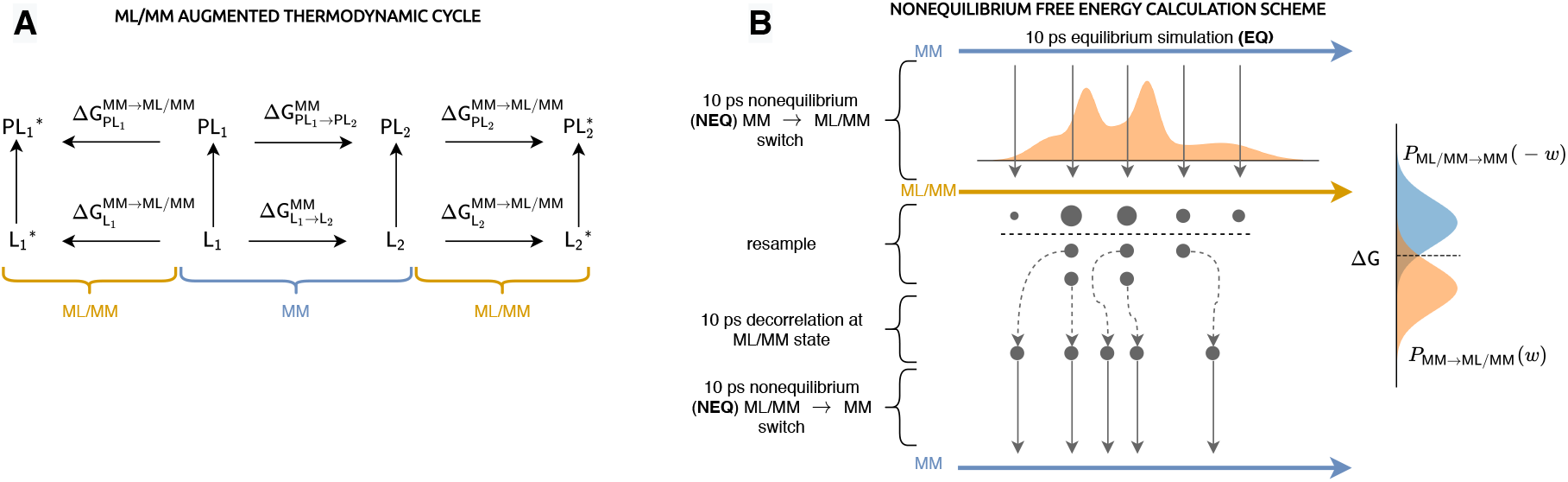
A simple nonequilibrium switching scheme can eficiently correct standard MM alchemical free energy calculations to ML/MM accuracy. **(A)** Augmented thermodynamic cycle used to estimate ML/MM free energies. The blue-bracketed, four-state thermodynamic cycle represents a typical MM relative free energy calculation where 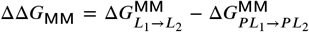. The orange bracketed thermodynamic augmentations represent respective ML/MM hybrid states such that 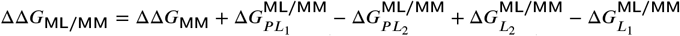. **(B)** Illustration of the nonequilibrium switching perturbation approach to estimating free energy corrections. The blue MM and orange ML/MM arrows represent equilibria at respective thermodynamic states. First, *N* configurations are sampled from equilibrium at the MM state and short MM→ML/MM nonequilibrium (NEQ) trajectories are generated and the dimensionless work *w*_MM→ML/MM_ recorded. Subsequently, the last configuration of each NEQ trajectory is resampled (with replacement) with probability proportional to 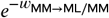. Resampled configurations are decorrelated with 10 ps of ML/MM equilibrium molecular dynamics before backward ML/MM→MM nonequilibrium trajectories are generated and the corresponding dimensionless work 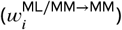 stored. The Bennett acceptance ratio (BAR) [54–56] is used to estimate the free energy difference, which corresponds to the crossing of the true *p*(*w*_MM→ML/MM_) and *p*(−*w*_ML/MM→MM_) work distributions [55, 57].

## Nonequilibrium perturbations can eficiently compute MM to ML/MM corrections

Current implementations of ML potentials do not permit an entire alchemical free energy calculation to be carried out with hybrid ML/MM potentials in a practical timescale. Instead, we aim for an approach that post-processes traditional MM alchemical free energy calculations [5, 12, 58]—such as the relative free energy calculation used here—to compute a correction Δ*G*^MM→ML/MM^ to the free energy of binding. Alchemical free energy calculations are now both routine and efficient, and available in a wide variety of software packages, often with GPU acceleration [6, 45, 50, 59–65].

While it may be tempting to simply sample from equilibrium MM states where the ligand is in complex or solution and estimate Δ*G*^MM→ML/MM^ based on the instantaneous dimensionless work *w*[*X*] ≡ *β*[*U*_ML/MM_(*X*) − *U*_MM_(*X*)], unless the MM models have been specifically parameterized to minimize the variance in *w* [41], small differences in the equilibrium valence degrees of freedom ensure that the variance of this weight is so large as to make this approach impractical (Figure 4).

**Figure 4.**
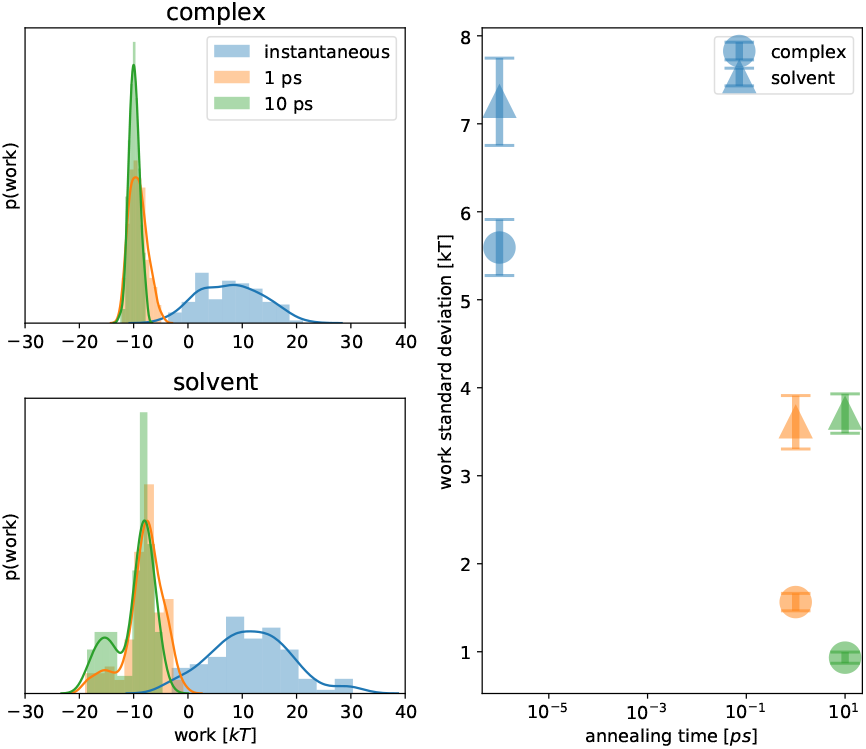
Short 10 ps nonequilibrium trajectories are suficient to reliably estimate MM→ML/MM free energy corrections. *Left*: Forward nonequilibrium work distribution (MM→ML/MM) for each of three switching times for both complex and solvated ligand phases for a representative ligand (**1**). The evolution of the work distributions demonstrates a reduction in variance and converge to more negative work values upon longer annealing time [69]. *Right*: Standard deviations of work distributions with respect to nonequilibrium protocol length for complex and solvent phases and bootstrapped 68% CIs. The complex phase has consistently lower work standard deviations, likely due comparatively lower ligand entropy than in the solvent phase.

As an alternative, we propose a convenient approach that uses short, nonequilibrium (NEQ) simulations to reduce the variance sufficiently to enable practical free energy estimates. Importantly, we aim to avoid two pitfalls: First, we aim to avoid the significant bias that arises in attempting to estimate free energy differences from short, unidirectional MM→ML/MM nonequilibrium switching trajectories [66], as well as the costly long nonequilibrium trajectories that would be required to minimize that bias; instead, we aim to use bidirectional protocols (MM→ML/MM and ML/MM→MM) and the optimal Bennett acceptance ratio estimator, which minimizes this bias [54, 55, 66]. Second, we aim to minimize the amount of simulation data that must be generated from the ML/MM state since it is so slow to sample from.

We construct an alchemical protocol that connects the easily-sampled MM thermodynamic states to the ML/MM thermodynamic states (in solvent and complex) via a linear interpolation of the potential (geometric interpolation of the sampled probability density function [67]) wherein the potential takes the form

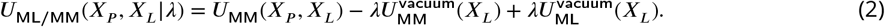

Here, the alchemical parameter *λ* ∈ [0, 1] interpolates between the MM and hybrid ML/MM endstates, 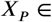 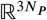 corresponds to the configuration of the receptor (and all other environment) atoms, and 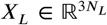 corresponds to the configuration of the ligand atoms. The NEQ free energy correction can be computed using four sequential steps, each performed independently for the solvated phase and the complex phase:

1. Extract *N iid* equilibrium samples from each MM thermodynamic state (ligand in complex or solvent) and perform MM→ML/MM nonequilibrium switching (NEQ) simulations for a fixed trajectory length *T* using the alchemical potential Eq. 2 with *λ* = *t*/*T*, recording the dimensionless protocol work *w*_MM→ML/MM_ [68].
2. Resample the final snapshots from each trajectory (with replacement) using the weight 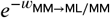 to generate an ensemble of *N* snapshots sampled from equilibrium for the ML/MM state.
3. For each resampled configuration, perform a short (10 ps) MD simulation with the ML/MM potential to decorrelate the resampled configurations.
4. For each of these ML/MM configurations, perform NEQ switching with the time-reversed protocol (ML/MM→MM) and record the dimensionless work *w*_ML/MM→MM_.

The Bennett acceptance ratio (BAR) [54–56] is then used to estimate the free energy correction Δ*G*^MM→ML/MM^ to the absolute free energy of the MM endstate for each phase (complex, solvent) of the alchemical free energy calculation.

We stress that resampling after forward NEQ switching (step 2) is a critically important part of the procedure. Once forward NEQ trajectories are collected, each final configuration is approximately Boltzmann distributed with respect to the ML/MM thermodynamic state as 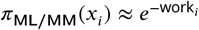 [70]. Omitting the resampling step and simply retaining all of the conformations generated by the forward NEQ switch step would not recover an approximately Boltzmann-distributed sample size. Omitting the resampling step would be particularly problematic in the solvent phase—where the forward work distributions generally span several *k_B_T* (see Fig. S.I.1)—since ligand conformations that are exponentially disfavored at the ML/MM state (compared to the MM state) would undergo backward NEQ switching, rendering prohibitively biased backward work distribution and free energy estimates. The resampling step is followed by a short equilibration (step 3) to decorrelate configurations that are resampled multiple times and recover from any collapse in effective sample size that occured during the resampling step.

An analysis of the unidirectional work distribution for several nonequilibrium protocol lengths (sometimes referred to as “annealing times” in the annealed importance sampling (AIS) literature [71]) suggests that 10 ps switching times are sufficient to produce useful free energy estimates (Figure 4). Indeed, performing the bidirectional switching scheme for several ligands confirms that the forward and backward work distributions overlap and BAR can produce useful estimates of the free energy corrections (Figure 5). Notably, solvent phase NEQ perturbations consistently yield higher work variances (Figure S.I.1) in their work distributions than their complex-phase counterparts, presumably indicative of the conformationally constrained nature of bound (but not solvated) ligands.

**Figure 5.**
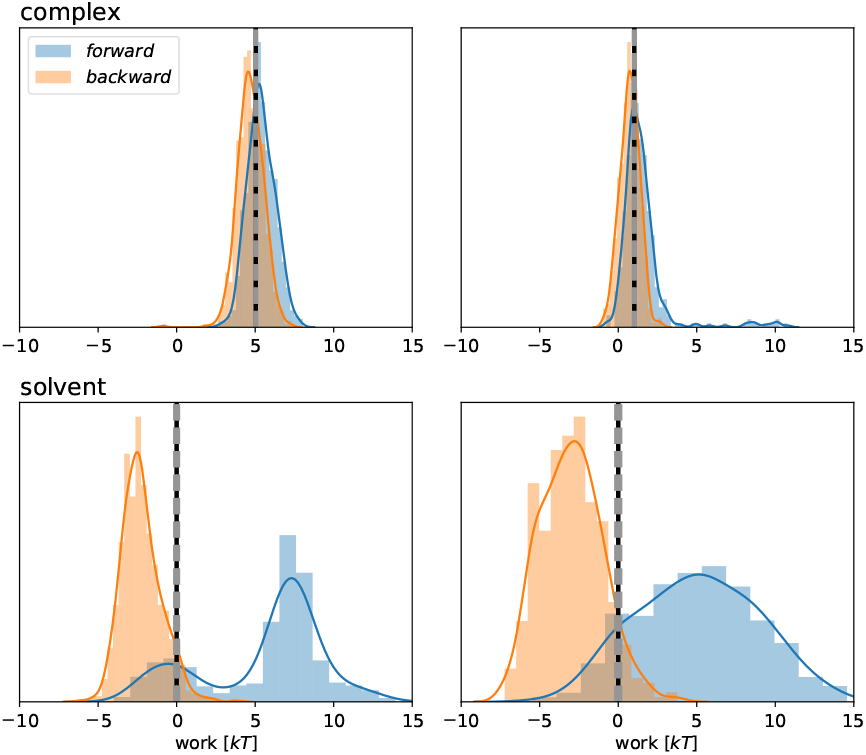
Bidirectional work distributions from nonequilibrium switching show sufficient overlap to compute precise free energy corrections from MM to ML/MM in both solvent and complex phases. Forward (blue) and negative backward (orange) work distributions from 10 ps nonequilibrium switching trajectories for MM to ML/MM perturbations are shown for complex (top) and solvent (bottom) phases for ligands 1 (left) and 14 (right). The Bennett acceptance ratio (BAR) estimates of the free energies and the uncertainty thereof are shown as vertical black lines and vertical gray dotted lines, respectively.

## ML/MM significantly improves accuracy on a kinase:inhibitor benchmark

We applied this nonequilibrium switching free energy correction scheme to a benchmark set of a well-studied congeneric series of inhibitors for non-receptor tyrosine-protein kinase (Tyk2) from the Schrödinger JACS benchmark set (Figure 2) [45]. This benchmark set is challenging for both commercial force fields (OPLS2.1 achieves a ΔG RMSE of 0.93±0.12 kcal mol^−1^ [45]) and public force fields (GAFF 1.8 achieves a ΔG RMSE 1.13 kcal/mol^−1^ and ΔΔG RMSE of 1.27 kcal/mol [50]). We consider a purely MM binding free energy baseline by using the ANI-2x [49] ML model (parameterized from DFT *ω*B97X/6-31G*) with AMBER14SB [47], TIP3P [48], and the Open Force Field Initiative OpenFF 1.0.0 (“Parsley”) small molecule force field [46].

The OpenFF 1.0.0 (“Parsley”) MM free energy calculations, shown in Figure 6 (**A**) and (**C**), achieve an accuracy that is statistically indistinguishable from other public and commercial MM force field benchmarks in terms of both root-mean squared error (RMSE; OPLS2.1 [45] and GAFF 1.8 [50]) and mean unsigned error (MUE; OPLS3.1, GAFF 2.1, and CGenFF 4.1 [64]). When the MM free energy calculation is corrected to ML/MM level of theory, Figure 6 (**B**) and (**D**), we recover experimental free energies with an RMSE of 0.47 [95% CI: 0.30, 0.67] kcal mol^−1^, a large and statistically significant improvement from MM (RMSE 0.97 [95% CI: 0.70, 1.22] kcal mol^−1^). Due to the naïve formulation of the ML/MM potential, this improvement in the experimental agreement can only be a consequence of an improved intramolecular potential for the ligands in this system. The particular formulation was chosen as it allows for rapid calculation of the per-ligand corrections to be performed *post hoc*. More advanced definitions of the ML/MM potential may lead to further improvements, by modelling additional interactions with higher levels of theory, however this would require concerted efforts to implement efficient interoperability between MM and ML packages—an area that requires further work.

**Figure 6.**
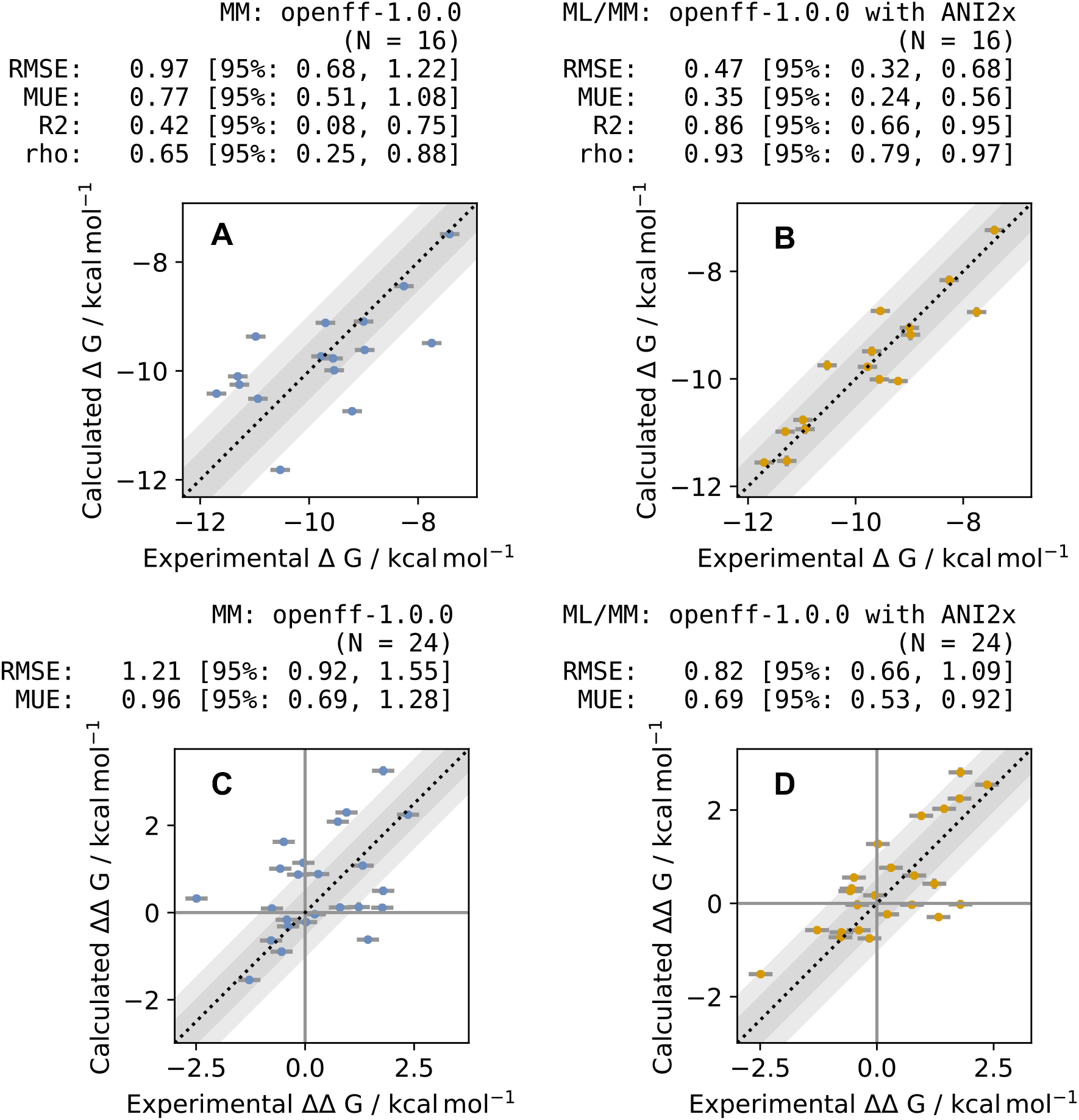
ML/MM free energy calculations show significant improvement over MM in reproducing absolute and relative Tyk2 inhibitor binding free energies. **(A)** Absolute binding free energies for the MM small molecule OpenFF 1.0.0 (“Parsley”) force field used with AMBER14SB and TIP3P water computed from relative free energy calculations estimated using perses 0.7.1 [http://github.com/choderalab/perses] and the maximum-likelihood estimator [72] to integrate estimates from redundant transformations in the relative alchemical transformation network. The same redundant network of relative alchemical transformations used in [45] was used here. **(B)** Absolute free energies (ΔG) corrected to ML/MM (using ANI-2x [49] for the ML model) using the nonequilibrium correction scheme depicted in Figure 3. **(C)** Relative MM binding free energies (ΔΔG) for computed relative free energy transformation edges, with correction using MLE. **(D)** Relative ML/MM binding free energies obtained from differences in the corrected absolute binding free energy estimates (top right). Blue scatter points are MM results, and orange are ML/MM results. Dark and light grey shaded regions indicate the region of ±0.5 and ±1.0 kcal mol^−1^ error respectively. Vertical error bars (which appear smaller than the symbols) show one standard deviation in the free energy, calculated by MBAR, while the experimental error bar of 0.18 kcal mol^−1^ is used [51]. Statistical analysis was performed using the Arsenic package [http://github.com/openforcefield/arsenic], with 95% confidence intervals calculated by bootstrap analysis. For all plots, an additive constant was added to all computed values, such that the mean computed value is equal to the mean experimental value, such as to minimise the RMSE as in [45].

The nature of these corrections is somewhat surprising: All MM→ML/MM corrections are positive, disfavorin binding. There is a notable trend in the magnitude of the correction, as illustrated by Figure 7. The smallest Δ*G*^MM→ML/MM^ corrections, on the order of 0.5–1.5 kcal mol^−1^ are the conformationally-strained cyclopropane moieties. Aliphatic groups with more conformational degrees of freedom, such as larger rings and acyclic groups, show larger corrections, with some of the largest MM→ML/MM corrections containing functional groups that will conjugate with the amide group. The subtleties of the electronics of these conjugated molecules are unlikely to be captured by MM forcefields, particularly small molecule torsion parameters [26].

**Figure 7.**
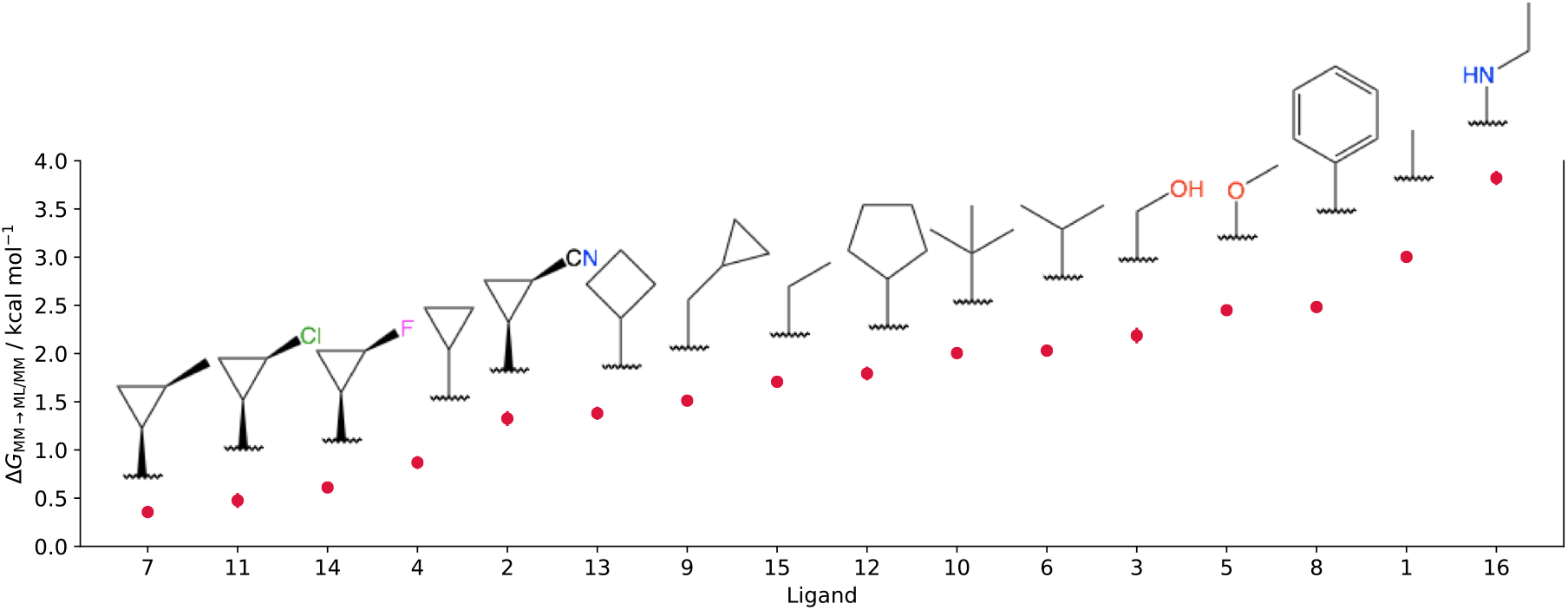
ML/MM corrections to MM binding free energies can be up to 4 kcal mol^−1^ in magnitude. The signed Δ*G*^MM→ML/MM^ corrections for each ligand (with R-group shown) are shown, ordered from least positive (slightly disfavoring binding) to most positive (strongly disfavoring binding).

## Improved torsion energetics appear to drive accuracy improvements

It is well-appreciated that general small molecule MM force fields often fail to accurately describe torsion energy profiles observed with higher-level quantum chemical calculations [73, 74], a phenomenon driven by the significant effect substituents can have on torsion profiles via electronic effects [26, 75]. To overcome this limitation, many MM force fields recommend refitting torsion potentials directly to quantum chemical calculations for individual molecules in a bespoke manner [1, 27].

It is plausible that the improved accuracy demonstrated by ML/MM in Figure 6 arises primarily from improved modeling of torsion energetics or torsion-torsion coupling. To investigate this, we examined the torsion probability density functions for conformations sampled by the ligand in solvent and in complex. Figure 8 depicts the 1D torsion (top) and 2D torsion-torsion (bottom) probability density functions for ligand **1**, focusing on the torsions associated with the amide linker to the substituted R-groups in the ligand series. The ML/MM potential samples a notably more peaked distribution with significantly perturbed equilibrium rotamer probabilities in solvent, and a tighter shifted torsion range in complex (Figure 8, top). 2D couplings are also surprisingly different between MM and ML/MM in complex.

**Figure 8.**
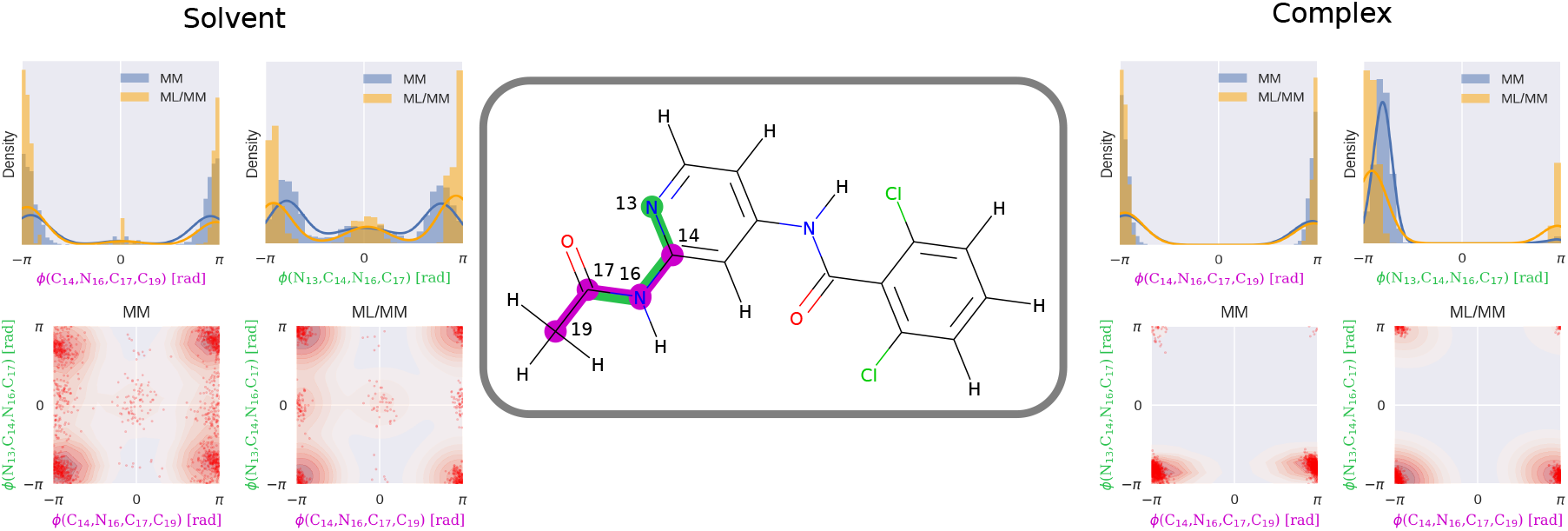
Torsion profiles and couplings differ between MM and hybrid ML/MM models. 1D and 2D torsion profiles of coupled amide torsions connecting the substituent R-group for ligand **1** (*center*) are shown. *Top row:* 1D torsion profiles for bonds highlighted purple and green for MM (blue) and MM/ML (orange) solvent and for complex. *Bottom row:* 2D torsion-torsion profiles for both solvent and complex are shown for both MM and ML/MM ensembles using a bivariate kernel density estimate, with red scatter points indicating observed samples.

## Discussion

In this work, we demonstrated that a hybrid ML/MM model of a challenging, pharmaceutically-relevant benchmark receptor:ligand system dramatically outperforms the both commercially and publicly-available MM force fields in its ability to recover experimental binding free energies of a congeneric series of inhibitors. The ability to halve the current state-of-the-art free energy uncertainty to ~0.5 kcal mol^−1^ using a simple post-processing procedure along with publicly-available software and ML models is promising. In particular, it suggests that there is significant potential yet to predict ligand:target binding affinities for prospective drug design campaigns.

### The NEQ protocol suggests the potential for further efficiency improvements

Among the insights made in the course of this study, we make special note of the NEQ procedure and its prospect for optimization. It is clear from Figures 4, 5, and the Supporting Information that a fixed-length NEQ protocol may indeed be wasteful in the complex phase considering the consistency of high work distribution overlap and BAR precision; in fact, exponential averaging results for all of the forward complex phase NEQ protocols yield free energies within 0.1 kcal mol^−1^ of the calculated BAR free energy correction. Indeed, reallocating the effort of conducting an ML/MM equilibration and backward protocol from complex to the solvent NEQ protocol might afford a more robust, lower variance free energy correction.

### ML/MM and MM torsion distribution discrepancies prompt further investigation

The substantial differences between torsion profiles from the ML/MM and MM models in Fig. 8 suggests that the significant free energy corrections afforded by the ML/MM model may be largely a consequence of poorly parameterized torsions in the MM model, or perhaps the existence of nonneglibible torsion couplings. Further experiments could distinguish between these scenarios by refitting torsion profiles or reweighting using only 1D or 2D torsion profiles rather than using the complete replacement of ligand intra molecular energetics as was considered here.

The fact that the most pronounced discrepancies in torsion profiles for both solvent and complex phases were observed about amide functional groups is particularly notable. If it is indeed the case that current MM force fields fail to recover appropriate conformational energetics of amides, then there is certainly a potential for free energy accuracy improvement in a large subset of druglike molecules, especially among protease inhibitors, which are characterized by several amide groups to mimic peptide backbones.

### Binding free energy improvements afforded by ML/MM show promise for other systems

In this study, we were fortuitous in that the OpenFF 1.0.0 provided heavy-tailed phase-space distributions, particular in the solvent phase (see Figure 8), that overlapped sufficiently with the ML/MM model. Had this not been the case, it is likely that the NEQ correction procedure would have failed to recover free energies with sufficient precision at the annealing times employed in this study. Further study will indicate whether other MM force fields—including the GAFF force field [13, 14] and more recent iterations of the OpenFF force fields—generally provide sufficient phase-space overlap with ML models for ML/MM corrections to remain computationally convenient and accurate with respect to experiment.

While these results illustrate the notable improvement to relative free energy calculations for this Tyk2 protein:ligand system, more extensive studies will be needed to determine how robustly this accuracy improvement manifests for a broad range of congeneric series. While the current implementation involves post-processing of pre-generated MM data, the implementation of the method could be improved by integrating ML potential models such as ANI into extensible simulation packages such as OpenMM [76], perhaps via a plugin architecture. Improved interoperability would increase ease of adoption for computational efforts in drug design projects. The definition of the hybrid ML/MM potential could be improved through expanding the terms in the system that are computed with ML by using ML methods such as AIMNet [77], SchNet [78], PhysNet [79], or AP-Net [80] that allow for decomposition of electrostatics and long-range dispersion from short-term valence energies.

### Machine learning will likely permeate all aspects of alchemical free energy calculations

More broadly, machine learning will play various roles in all aspects of alchemical free energy calculations. As more calculations are performed, machine learning models (such as graph convolutional or message passing networks [81]) will undoubtedly be used to learn the *difficulty* (statistical efficiency) of relative transformations in a manner that can be used to design optimal transformation networks [72] or optimal alchemical protocols. Scheen et al. [82] recently demonstrated how ligand-based ML models can be used to correct MM alchemical free energy calculations based on experimental training data, applying it to hydration free energy computation. Ghanakota et al. [83] also demonstrated how ligand-based ML models could be trained to learn more expensive free energy calculations to permit evaluation of large compound spaces with free energy accuracy. These few applications are just the beginning of how machine learning will transform physical modeling in the biosciences.

## Code and data availability

• Input files and setup scripts: https://github.com/choderalab/qmlify

## Author Contributions

Conceptualization: JDC, DAR, HEBM, JF, and MW; Methodology: JDC, HBM, DAR, JF, and MW; Software: PBG, DAR, HBM, JF, and MW; Investigation: DAR, HEBM, JF, and MW; Writing–Original Draft: DAR, JDC, HBM, JF, and MW; Writing–Review&Editing: AER, OI and JDC; Funding Acquisition: JDC; Resources: JDC; Supervision: JDC, AER, and OI.

## Acknowledgments

DAR acknowledges support from the Tri-Institutaional PhD Program in Chemical Biology and the Sloan Kettering Institute. HEBM acknowledges support from a Molecular Sciences Software Institute Investment Fellowship and Relay Therapeutics. JF acknowledges support from NSF CHE-1738979 and the Sloan Kettering Institute. MW acknowledges support from a FWF Erwin Schrödinger Postdoctoral Fellowship J 4245-N28. JDC acknowledges support from NIH grant P30 CA008748, NIH grant R01 GM121505, NIH grant R01 GM132386, and the Sloan Kettering Institute. OI acknowledges support from NSF CHE-1802789 and Carnegie Mellon University. AER acknowledges support from NSF CHE-1802831

The authors thank Christopher Rowley (ORCID: 0000-0002-0205-952X) for sharing early work and discussions that inspired this study; Christopher I. Bayly (ORCID: 0000-0001-9145-6457) and David L. Mobley (ORCID: 0000-0002-1083-5533) for sharing insights on nonequilibrium free energy calculations; Gianni de Fabritiis (ORCID: 0000-0003-3913-4877) for discussions and inspirational work in motivating the utility of hybrid ML/MM models; Peter K. Eastman (ORCID: 0000-0002-9566-9684) for providing extensive support for OpenMM and implementing numerous features of use in this work; and the scientists and software scientists from the Open Force Field Initiative [http://openforcefield.org/members] and Consortium [http://openforcefield.org/consortium] for their contributions to both the science behind OpenFF 1.0.0 and the highly usable software infrastructure that made this work possible.

The authors are extremely grateful to OpenEye Scientific for granting an academic license for use of the OpenEye Toolkit for this work.

## Disclosures

JDC is a current member of the Scientific Advisory Board of OpenEye Scientific Software and a consultant to Foresite Laboratories. The Chodera laboratory receives or has received funding from multiple sources, including the National Institutes of Health, the National Science Foundation, the Parker Institute for Cancer Immunotherapy, Relay Therapeutics, Entasis Therapeutics, Silicon Therapeutics, EMD Serono (Merck KGaA), AstraZeneca, Vir Biotechnology, Bayer, XtalPi, the Molecular Sciences Software Institute, the Starr Cancer Consortium, the Open Force Field Consortium, Cycle for Survival, a Louis V. Gerstner Young Investigator Award, and the Sloan Kettering Institute. A complete funding history for the Chodera lab can be found at http://choderalab.org/funding AER is a current member of the Science Advisory Board of Schrödinger Co. and receives funding from Genentech Co.

## Detailed methods

### MM Relative Free Energy Calculations

MM relative free energy calculations were performed using the open-source software Perses 0.7.1 [http://github/choderalab/perses]. The set of 24 pairwise comparisons for the set of 16 ligands were performed, as specified in [45]. Simulations were performed for both the bound-complex phase and solvent phase to afford relative binding free energies using OpenMM 7.4.2 [76]. Protein and ligand files were adapted from [50] and are available as part of the openmm-forcefields 0.7.1 package [https://github.com/openmm/openmmforcefields]. Simulations were performed with AMBER14SB [47] protein forcefield and the OpenFF 1.0.0 “Parsley” small molecule forcefield [46]. The system was solvated with a 9.0 Å padding using TIP3P water [48] and 150 mM NaCl [84]. Simulations were performed without hydrogen bond constraints using 4 amu hydrogen massses following mass repartitioning. A timestep of 2 fs with a 1 ps^−1^ collision rate at 300 K was performed using a BAOAB Langevin integration scheme [85, 86], using an NPT ensemble at 1.0 atm sampled using a Monte Carlo barostat with molecular scaling. Nonbonded interactions were handled using a 9 Å cutoff using Particle Mesh Ewald (PME) with a tolerance of 2.5 × 10^−4^ and long-range dispersion corrections.

The alchemical perturbation was performed using a single topology protocol. The mapping protocol to generate the single hybrid ligand is performed using the maximum common substructure search (MCSS) algorithm from the OpenEye Toolkits 2020.0.4 to identify the common ‘core’ of the molecule. Atoms not common between the two molecules (not contained in the core) are included as ‘unique-old’ or ‘unique-new’ atoms. The interaction perturbation scheme is described in Figure S.I.2. Softcore steric potentials (see [87]) with an *α* parameter of 0.85.

11 equally-spaced *λ*-windows were used for these calculations, for both the solvent and the complex phases. 1000 cycles of 250 integration steps (2 fs timestep) were performed, resulting in a total of 5 ns of sampling per *λ*-state (55 ns sampling per-phase, per-ligand pair). All-to-all Hamiltonian replica exchange was attempted every cycle. MBAR was performed on decorrelated replica exchange samples to recover MM relative free energies and associated uncertainties. The maximum likelihood estimator (MLE) DiffNet [72] was used to compute absolute binding free energies for the set of ligands, and shifted using a single experimental value. The aforementioned calculations are self-contained operations in the [https://github.com/openforcefield/arsenic] as a graph representation (nodes correspond to ligands and edges correspond to relative alchemical transformations). Results are shown in Figure 6 (**A** and **C**) and were used as the basis for the MM→MM/ML corrected energies, described in the following.

### Bidirectional nonequilibrium switching and ML/MM free energy corrections

The bidirectional nonequilibrium protocol was parameterized in accordance with Eq. 2 with 5000 annealing steps. MM model and simulation parameters are consistent with those described above.

The forward NEQ procedure is as follows for each phase:

#### Algorithm 1: Nonequilibrium (NEQ) switching protocol

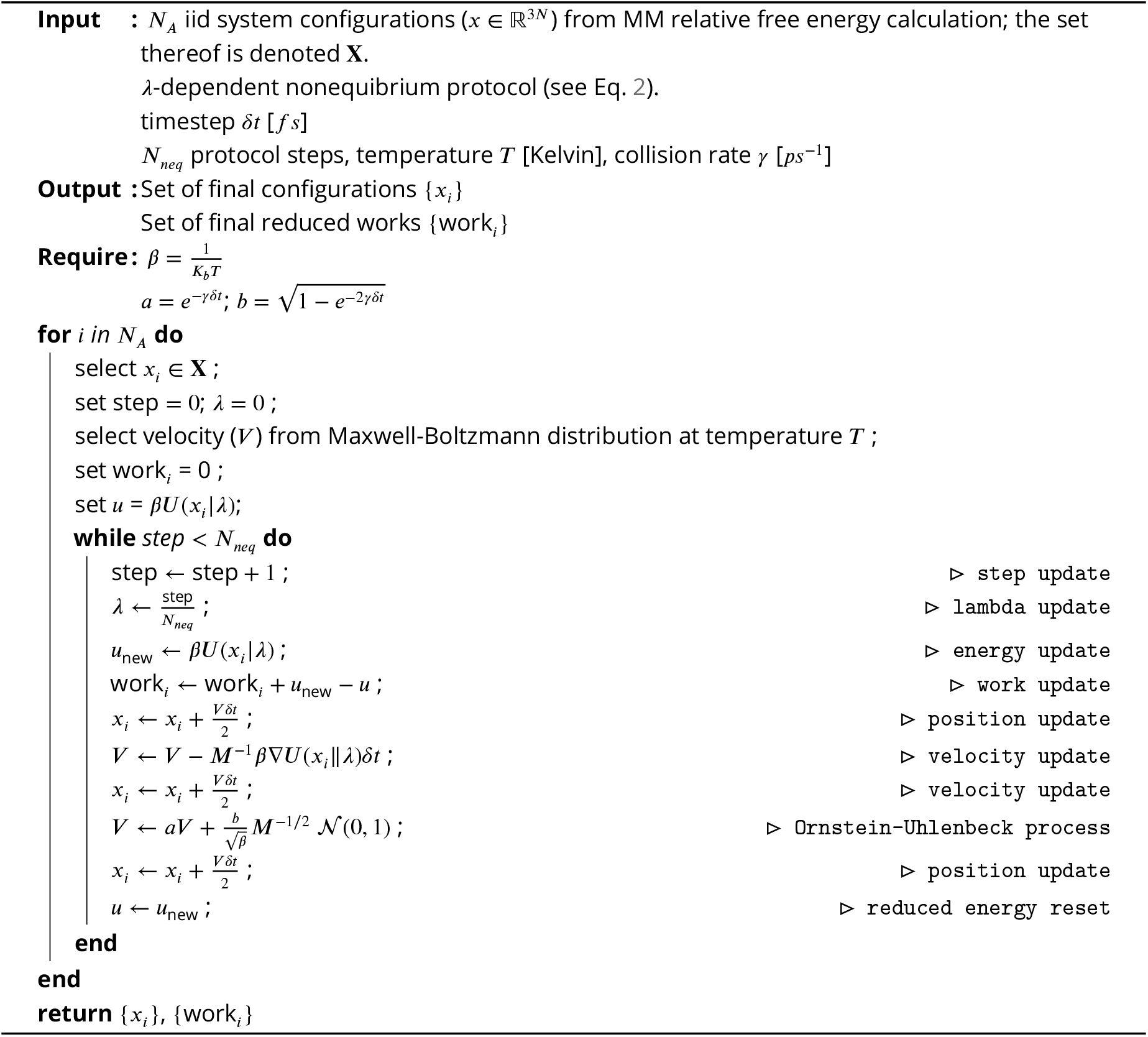

To conduct decorrelation and equilibration at the ML/MM thermodynamic states, {*x_i_*} returned from Algorithm 1 is resampled *N_A_* times (with replacement) w.r.t. 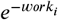) from {work_*i*_}. The algorithm is performed again (in this case, setting *N_neq_* to 5000) whilst maintaining *λ* = 1; in this case, work_*i*_ is identically 0. Subsequently, Algorithm 1 is conducted in the backward direction (i.e. 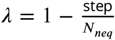 in the “lambda update” step).

The work sets {work_*i*,forward_} from Algorithm 1 and {work_*i,*backward_} from the aforementioned modification were then passed to the BAR estimator [https://github.com/choderalab/pymbar/blob/master/pymbar/bar.py] to maximize the likelihood of the Δ*G_ML/MM_*. To correct the MM free energy calculations, the complex/solvent 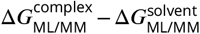 differences were added in-place to the MM ligand network nodes, and absolute binding free energies were recomputed with DiffNett [72].

For both the complex and solvent phase, 100 decorrelated equilibrium configurations (*N_A_* = 100) were extracted from replica exchange checkpoint files from each relative calculation edge. This treatment resulted in duplicate and independent repeats of ensemble ML/MM bidirectional switching. This resulted in 200 to 700 independent forward/backward work samples for each phase for the 16 ligands, depending on the degree of each ligand in the relative free energy calculation network. The aggregated bidirectional work distributions for each ligand and each phase is shown in Figure S.I.1.

The forward, resampling, ML/MM endstate simulation (for 10 ps), and backward annealing procedures, described previously, were used to recover bidirectional work distributions along with the BAR-estimated free energy correction.

The ligand configuration was extracted from the solvent and complex phases and modelled with the OpenFF 1.0.0 forcefield as used in the MM simulations; however the nonbonded (i.e. steric and electrostatic) interactions were treated as non-periodic without a cutoff. The ANI2x ligand model was computed using TorchANI package [https://github.com/aiqm/torchani] (ANI2x version 2.1.1)[88].

In order to propagate dynamics through the protocol, each velocity update steps are preceded by a force update step wherein the forces of the MM and ML vacuum ligand systems are computed, scaled appropriately by the value of *λ*, and added to the appropriate sub-block matrix of the full ligand-and-environment force matrix.

## Supplementary Information

**Appendix 0 Figure S.I.1.**
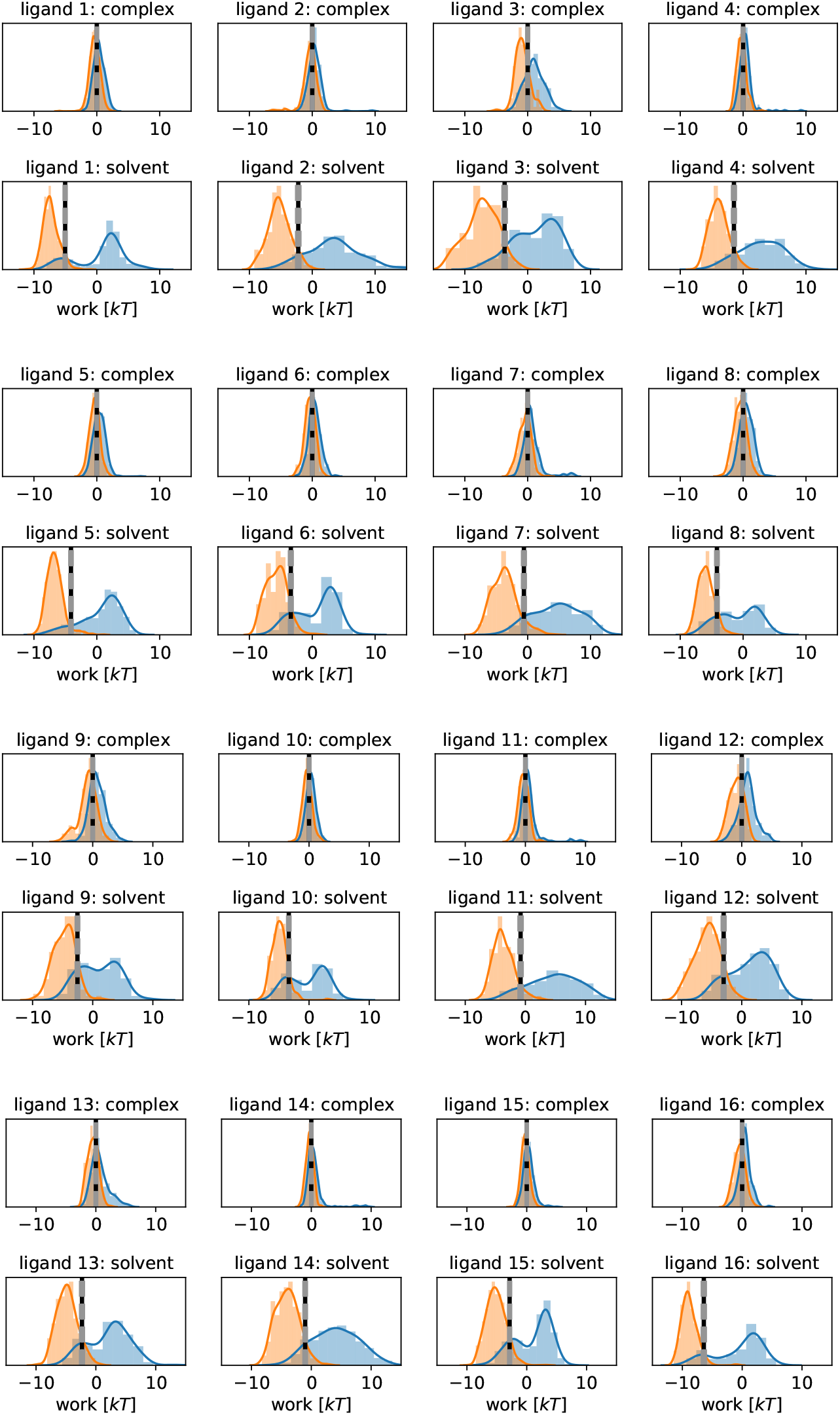
Bidirectional NEQ work distributions for Tyk2 congeneric inhibitor series. Forward work distributions (blue) and (negative) backward work distributions (orange) show sufficient overlap for a BAR free energy calculation. BAR Δ*G_MM→MM/ML_* are shown as vertical black lines, and uncertainties thereof are shown as gray, dotted lines. Both complex and solvent phases are shown.

**Appendix 0 Figure S.I.2.**
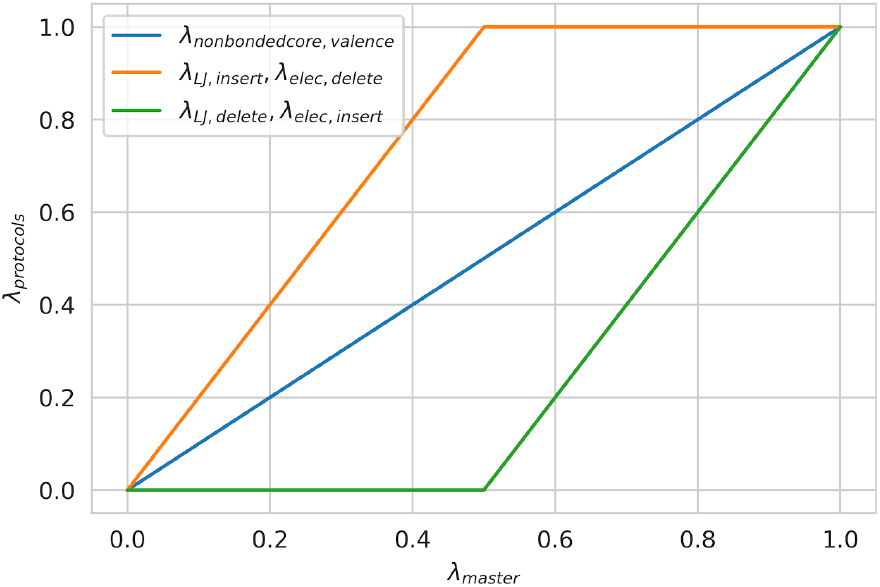
Perses relative free energy calculations default alchemical protocol. The interaction potentials are perturbed linearly in two stages: between *λ*: 0.0 → 0.5 the sterics of the unique-new atoms are turned on while the electrostatics of the unique-old atoms are turned off, followed by the turning on of the electrostatics of the unique-new atoms simultaneously with the steric terms of the unique-old atoms being turned off between *λ*: 0.5 → 1.0. The parameters of the core atoms are linearly perturbed from *λ*: 0. → 1.0.

## Notes

https://github.com/choderalab/qmlify

